# Tuning antibody stability and function by rational designs of framework mutations

**DOI:** 10.1101/2025.03.13.642419

**Authors:** Joseph C. F. Ng, Alicia Chenoweth, Maria Laura De Sciscio, Melanie Grandits, Anthony Cheung, Tooki Chu, Alexandra McCraw, Jitesh Chauhan, Yi Liu, Dongjun Guo, Semil Patel, Alice Kosmider, Daniela Iancu, Sophia N. Karagiannis, Franca Fraternali

## Abstract

Artificial intelligence and machine learning models have been developed to engineer antibodies for specific recognition of antigens, however these approaches often focus on the antibody complementarity determining region (CDR) whilst ignoring the immunoglobulin framework (FW) which provides structural rigidity and support for the flexible CDR loops. Here we present an integrated computational-experimental workflow, combining static structure analyses, molecular dynamics simulations and *in vitro* physicochemical and functional assays to generate rational designs of FW mutations for modulating antibody stability and activity. We first showed that recent antibody-specific language models lacked insights in FW mutagenesis, in comparison to approaches which utilised antibody structure information. Using the widely-used breast cancer therapeutic trastuzumab as a use case, we designed stabilising mutants which were distal to the CDR and preserved the antibody’s functionality to engage its cognate antigen (HER2) and induce antibody-dependent cellular cytotoxicity (ADCC). Interestingly, guided by local backbone motions predicted using molecular dynamics simulations, we designed a FW mutation on the trastuzumab light chain which retained antigen-binding effects but lost Fab-mediated and Fc-mediated effector functions. This highlighted effects of FW on immunological functions engendered in distal areas of the antibody, and the importance to consider attributes other than binding affinity when assessing antibody function. Our approach incorporates interdomain dynamics and distal effects between FW and the Fc domains, expands the scope of antibody engineering beyond the CDR, and underscores the importance of a holistic perspective that considers the entire antibody structure as a whole in optimising antibody stability, developability and function.

## Introduction

Antibodies are immunoglobulin molecules that recognise and bind to specific antigens, as a hallmark of the adaptive immune response to clear any invading pathogens, or to aberrantly expressed molecules in contexts such as cancers^1–4^. Recent advances in machine learning (ML) and artificial intelligence (AI) approaches can be applied to study how antibody responses are mounted *in vivo*^5–7^, and to design monoclonal antibodies (mAbs) as therapeutics that engage specifically to different clinically relevant targets^8–10^. In particular, protein language models have been applied to evolve antibodies *in silico*, towards better, more stable antigen binders^11–13^. The advancement of AI approaches also motivated high-throughput profiling of random antibody libraries, to generate experimental data for the training of next-generation antibody ML models^12,14^. The focus of such efforts has typically been in the fragment antigen-binding (Fab) region of the antibody, specifically the complementarity determining regions (CDRs), which are the major contributor to antigen binding (**Fig. 1a**): the CDR loops form a “paratope” of specific three-dimensional shape, which determines its antigen specificity^15,16^. The third CDR loop on the antibody heavy-chain, hereafter CDRH3, is a known major contributor to such specificity^12,16–18^. For example, Chinery et al.^14^ generated over 500,000 CDRH3 variants of the breast cancer therapeutic trastuzumab, to benchmark the performance of a variety of computational methods in predicting high-affinity antibodies from these randomly generated sequence libraries, and designing new antigen binders *de novo*. Furthermore, deep learning approaches identified CDR mutations with the potential to enhance the affinity and stability of therapeutic antibodies^12,13^.

**Figure 1.**
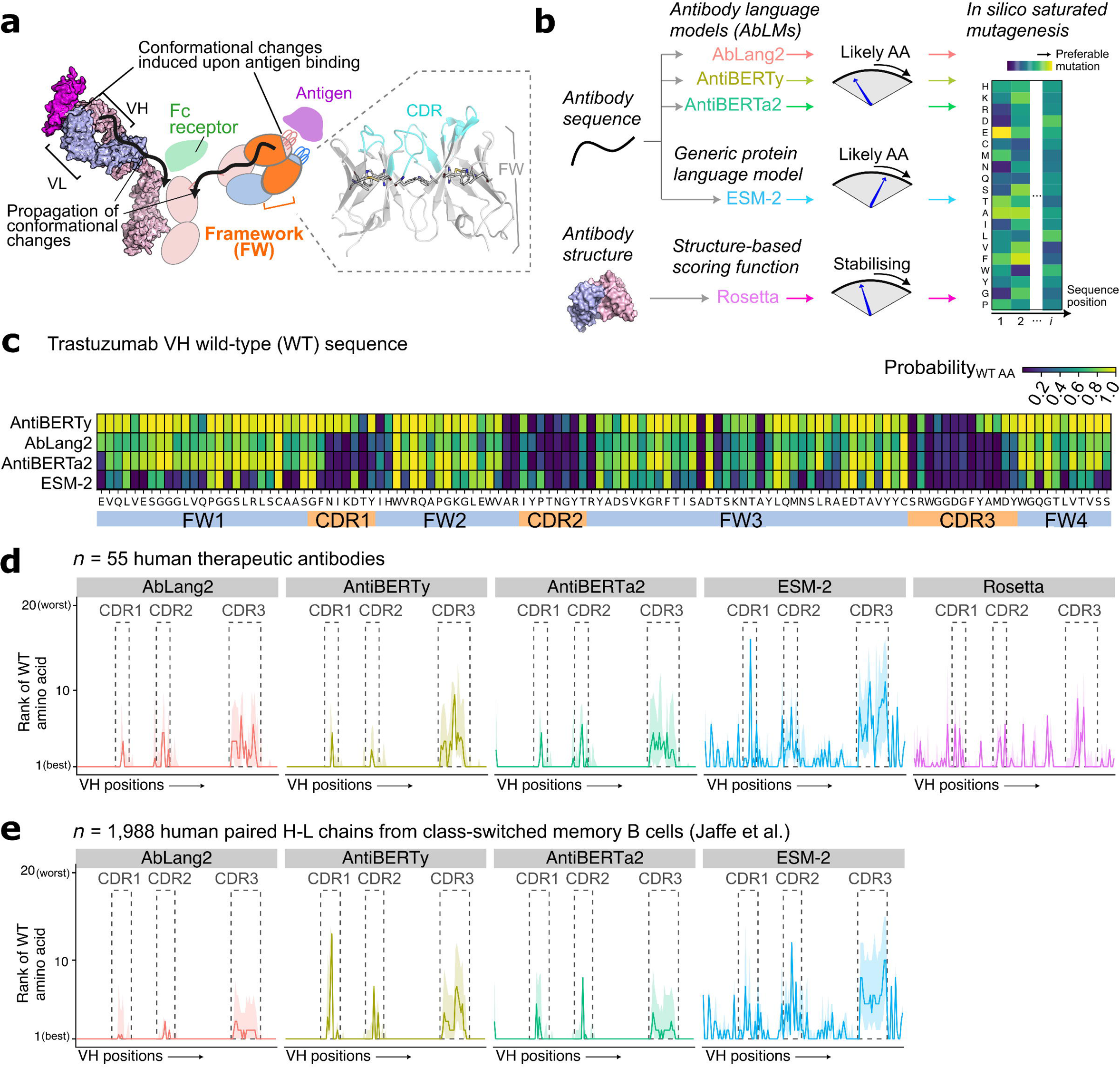
Antibody language models (AbLMs) offer limited insights for antibody framework (FW) mutagenesis. (a) Schematic of the role of FW region in antibody structure. (*left*) Antibody complementarity determining regions (CDR) undergo conformational changes upon antigen binding. These changes are thought to propagate through the FW to the rest of the molecule and can affect effector functions. (*right*) Illustration of CDR and FW on the antibody variable heavy (VH) and variable light (VL) region structure. Conserved positions in FW (ref. 53) are highlighted with stick representation. (b) Illustration of *in silico* saturated mutagenesis using AbLMs, a generic protein language model (ESM-2) and a structure-based approach (Rosetta). (c) Heatmap illustrating position-specific probabilities for the wild-type amino acid (columns) along the trastuzumab VH sequence using different language models (rows). FW and CDR regions are delimited below the heatmap. (d-e) Rank of the wild-type amino acid (vertical axis, 1 = best, 20 = worst) along the VH sequences (horizontal axis) of (d) *n* = 55 human therapeutic antibodies and (e) n = 1,988 paired H-L chains from class-switched memory B cells taken from Jaffe et al. (ref. 52), according to the approaches evaluated in (b). For (e) Rosetta was not attempted due to unavailability of experimentally determined structures for these sequences. Lines depict median and shaded areas depict the inter-quartile range.

However, the function of an antibody is determined not only by its binding affinity to the antigen, but also by a number of other factors. The term of antibody “developability” has been used to encompass a range of issues pertaining to the potential success of antibody designs, including stability, solubility and immunogenicity, amongst others^19–22^. Computational methods to predict these properties have been developed to streamline *in silico* antibody design^22–26^. In addition, the therapeutic effects of mAbs do not solely arise from antigen binding, but also from effector functions mediated by the fragment crystallisable (Fc) region, such as antibody-dependent cellular cytotoxicity (ADCC)^27–29^. These factors require consideration beyond the CDR regions and the antibody-antigen interface^27,30,31^. In antibodies, CDR regions can be found in both the heavy and the light chain variable domains, which comprised chiefly of an immunoglobulin fold “framework” (FW) (**Fig. 1a**). The FW region provides structural integrity to support the flexible CDRs^15,30–34^. Positioned between the antigen-binding CDR region and the antibody constant (C) region, the FW immunoglobulin folds in the heavy and light chains interact with distinct packing geometries, which are thought to mediate the propagation of conformational changes from the CDRs to the rest of the antibody upon antigen binding^17,35–40^. If we consider antibody structures under this holistic perspective^17,41,42^, exclusive engineering of the antigen-antibody interface may yield unintended consequences if these perturbations affect the FW and consequently the propagation of conformational changes throughout the antibody molecule^17,35,43–45^, which could result in adverse effects or a loss of functional efficacy. There is a paucity of detailed molecular understanding on how the FW confers such effects, and how FW mutations could disrupt both antigen binding and Fc-mediated effector functions. Conversely, FW mutations can also be harnessed to optimise these properties without directly altering the antigen-binding site, representing a promising avenue to subtly modulate antibody stability and function^18,31,33^.

Here, using the widely-used breast cancer therapeutic trastuzumab as a model system, we conducted iterative computational-experimental analyses to investigate mutations on both the trastuzumab heavy and light chains, exploring the potential of FW mutagenesis to tune antibody functionality. Trastuzumab is a well-established therapeutic specific to human epidermal growth factor receptor 2 (HER2), overexpressed in ∼15% of breast cancers, and is the first-line treatment to these cancers^46^. We highlighted the limitations of existing protein language models trained solely on antibody repertoire data (hereafter antibody language models, or AbLMs) for FW mutagenesis, and opted instead for a structure-based approach to predict stabilising FW mutations, and verified the stability, antigen binding and downstream functional effects of these mutants using *in vitro* assays. We employed *in silico* molecular dynamics (MD) simulations both to elucidate insights on how the designed mutants perturbed antibody structure, and also to suggest new mutants based on antibody local structural dynamics. We dissected the contribution of antibody FWs to antigen-binding and effector functions, fine-tuning antibody stability and function using carefully designed FW mutants.

## Results

### Antibody language models offer limited insights in FW mutagenesis

We first assessed different computational approaches in their usefulness to suggest mutations in antibody FW regions. In our comparisons we considered three types of approaches which are capable to generate numerical scores of mutational impact: (i) biological language models trained on generic protein sequences^47^; (ii) antibody language models^5,48,49^ (AbLMs), and (iii) structure-based approaches which evaluate mutational effects using a biophysical scoring function that considers different components of intramolecular forces^50,51^. In total we considered five computational approaches (**Fig. 1b**), inputting either the protein structure or sequence of trastuzumab variable heavy (VH) and variable light (VL) domains, and extracting the predicted effects of mutating every amino acid position to every other nineteen amino acids, i.e. performing an *in silico* saturated mutagenesis scan. **Fig. 1c** displayed the probability of the wild-type (WT) amino acid at each position along the trastuzumab VH sequence, according to the four language models evaluated in this analysis. We noticed that all tested language models were uncertain about amino acid identities in the CDR region, but more confident about the FW region. This was especially true for the AbLMs, which consistently gave a probability close to 1 for WT amino acids at the FW positions (**Fig. 1c**). In contrast, ESM-2, which was not trained specifically on antibody sequences, displayed a more varied pattern. For the VL sequence, similar patterns were observed, with the exception that AbLMs were more confident in the CDR loops than in the case of the VH domain (Supplementary Figure S1).

We repeated this evaluation over *n* = 55 human therapeutic antibodies with structural information, which reinforced the patterns seen for trastuzumab (**Fig. 1d**; Supplementary Figure S2); here, similar to ESM-2, structure-based predictions given by the Rosetta point mutant scan application were also more varied in the FW. This could either reflect the fact that the input sequences were well-established, manufacturable antibody therapeutics with little room for further optimisation in the FW, or that the AbLMs were over-specialising to antibodies whilst failed to generalise to mutations which were rarely seen, such as those in the FW region. We explored this by utilising a dataset of paired VH and VL sequences from sorted switched memory B cells from *n* = 3 healthy donors^52^, and observed a similar pattern (**Fig. 1e**; Supplementary Figure S3). This suggested that, despite recent interests in the field of antibody discovery due to their ability for efficient generative designs^12^, AbLMs focus mainly on CDRs in detriment of their capabilities to suggest new FW mutations that would enhance antibody stability.

### *In silico* structure-based screening identified VH S85N+R87T as a stabilising and functional trastuzumab variant

Based on the previous analysis, we opted to pursue *in silico* FW mutagenesis using a structure-based approach to evaluate the stabilising effects of mutations, by estimating free energy change (ΔΔ*G*) of the antibody upon mutation. Using the Rosetta molecular modelling software, we used the point mutant (pmut) scan application to predict ΔΔ*G* of every possible amino acid substitution along the trastuzumab VH and VL sequences. Most (94.19% in the VH and 94.20% in the VL) Rosetta-predicted mutations were destabilising (**Fig. 2a**), in line with the fact that trastuzumab is a well-used therapeutic antibody with VH and VL domains close to their optimal properties.

**Figure 2.**
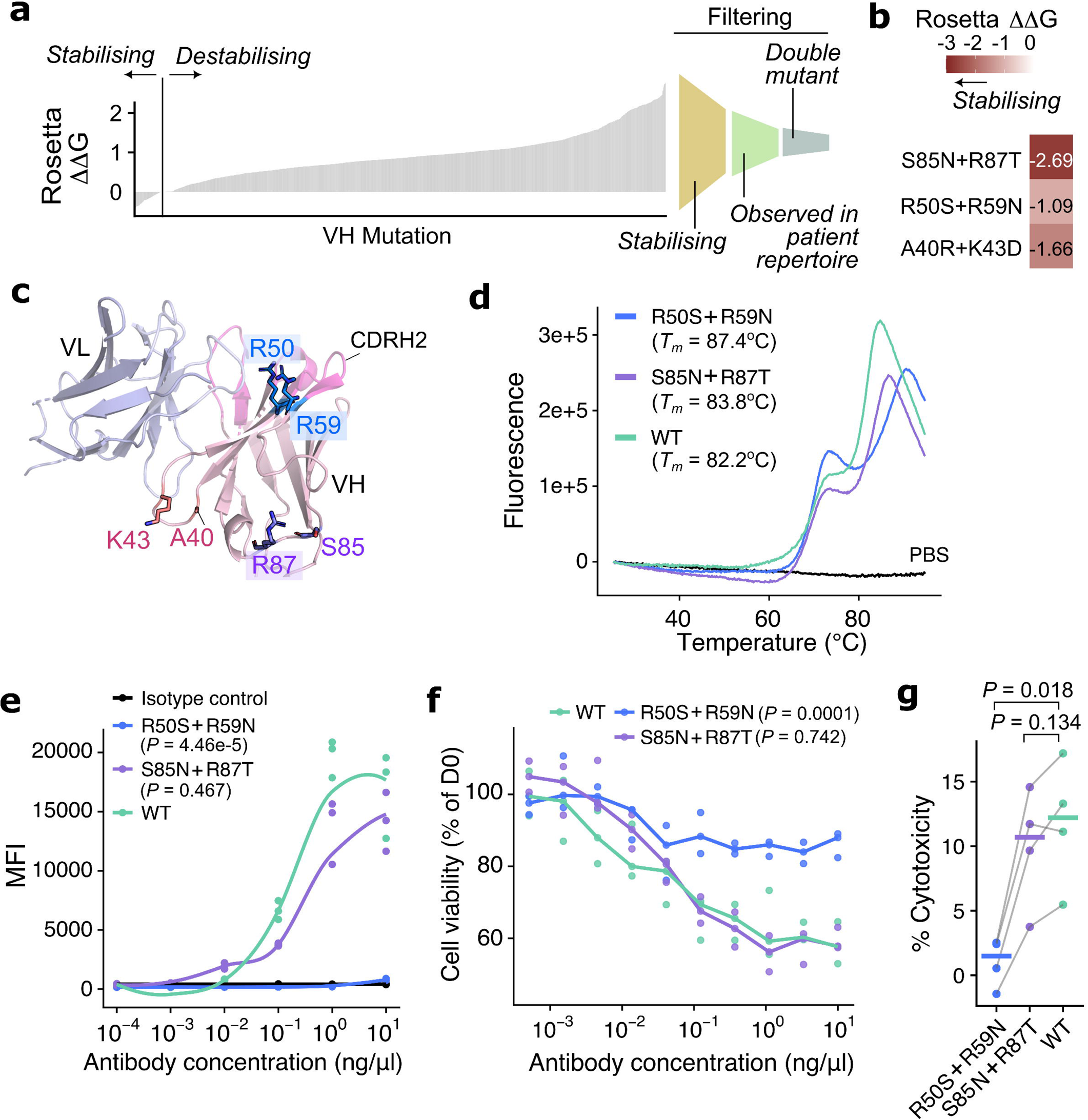
Structure-based designs of trastuzumab FW stabilising mutations. (a) (*left*) Bar-plot depicting Rosetta-predicted ΔΔG for every possible amino acid substitution in the trastuzumab VH structure, using PDB 7mn8 as input. (*right*) Filtering approach to select mutations for investigation. (b) Heatmap depicting Rosetta-predicted ΔΔG for the three double mutants selected for further characterisation. (c) Illustration of the positions of double mutants listed in (b) in the trastuzumab structure. (d) Melting curves of R50S+R59N and S85N+R87T with respect to the wild-type (WT), determined using differential scanning fluorimetry. PBS-only condition served as a negative control. The melting temperature (*T_m_*) of the constructs are indicated. (e) Binding of WT and mutants against a HER-2 expressing cancer cell line (SKBR3) determined using fluorescence-activated cell sorting (FACS). Statistical significance was evaluated using analysis of variance (ANOVA) models and *P*-values were obtained by post-hoc Dunnett’s test, comparing mutants to the WT. (f) Viability of SKBR3 cells *in vitro* in the presence of the antibody constructs. Cell viability was measured as a percentage of cell counts at Day 0 of the experiment. Statistical significance was evaluated using ANOVA followed by post-hoc Dunnett’s tests comparing mutants to WT. (g) Antibody-dependent cellular cytotoxicity (ADCC) effect of the antibody constructs, measured on SKBR3 cells (quantified using lactase dehydrogenase [LDH] release as a proxy) upon co-culturing with natural killer (NK) cells isolated from n = 4 healthy donors. P-values were quantified using paired t-tests and noted on the plot after false discovery rate correction.

To further filter for candidate FW mutations which optimise antibody stability, we focussed on the VH domain and applied the following criteria (**Fig. 2a**): first, we analysed the occurrence of these predicted mutations in antibody repertoires from breast tumour samples^2^. We reasoned that naturally occurring residues along the antibody sequence represent tolerable amino acids for stability and function. We observed that 50% of VH stabilising mutations were observed in >1% of the repertoire sequences (Supplementary Figure S4). Second, we performed an additional round of Rosetta mutational scanning for double VH mutants, in order to augment the chance to obtain variants with measurable effects on trastuzumab stability and function. We prioritised mutants with consistent Rosetta results between the single and the double mutant scans, and designed three stabilising VH double-mutants located at distinct sites within the FW region (**Fig. 2b**): A40R+K43D, R50S+R59N and S85N+R87T. R50 and R59 are situated at the apex of the β-fold (**Fig. 2c**). Despite being located at the base of the CDRH2 loop, these positions were classified as the FW region in the IMGT (international ImMunoGeneTics) numbering system^53^. A40R and K43D were each predicted as destabilising individually (Supplementary Table S1), but the combined effect of the double mutant was predicted to be stabilising (**Fig. 2b**), potentially due to compensatory effects of the two mutations on the surface charge of the antibody. S85 and R87 were both solvent-exposed positions at the opposite side to the CDR region (**Fig. 2c**).

We next verified the effects of these trastuzumab mutants using *in vitro* assays. We focussed on S85N+R87T and R50S+R59N, since the A40R+K43D double mutant had low production yield insufficient for further characterisation. We explored possible reasons for this subsequently in analysing our molecular dynamics (MD) simulation data (see below). For the remaining two variants, we determined their thermostability using differential scanning fluorimetry. We found that both mutants were stabilising compared to wild-type (WT), as indicated by a higher melting temperature (*T_m_*) (**Fig. 2d**), validating our *in silico* predictions.

We also assessed the capability of each antibody to bind its cognate antigen via fluorescence-activated cell sorting (FACS) against a HER2-expressing breast cancer cell line (SKBR3). We observed a significant reduction of binding for the R50S+R59N mutant (**Fig. 2e**). When we assessed Fab-mediated effects on cancer cell viability and Fc-mediated immune cell effects against cancer cells in antibody-dependent cellular cytotoxicity (ADCC) functional assays, we found that the R50S+R59N mutant also lost capability to inhibit SKBR3 cell proliferation (**Fig. 2f**) and to induce ADCC (**Fig. 2g**). The loss of functionality for R50S+R59N is consistent with their structural localisation (**Fig. 2c**), being in vicinity to the CDR region as predicted in our structural analysis.

To further elucidate our *in vitro* evidence from a detailed structural perspective, MD simulations were carried out on the designed trastuzumab variants as well as the WT antibody, in complex with HER2 (**Fig. 3a**). We generated and analysed MD trajectories of a total of 3 μs for each system (Supplementary Table S2). The overall conformational stability of the fragment antigen-binding (Fab) was retained for WT as well as the designed variants (Supplementary Figure S5), in agreement with the high *T_m_* measured experimentally. Analysing the WT trajectories, we observed that the loop containing A40 and K43 exhibited substantial flexibility (Supplementary Figure S6); the A40R+K43D mutant could potentially form a salt bridge to rigidify this loop and induce structural instabilities, which could be a possible reason for its low yield in our *in vitro* antibody production. As a proxy of the binding strengths between our trastuzumab variants and HER2, the distance between the centre of mass (COM) of the CDR region and COM of the HER2 juxtamembrane domain was monitored along the trajectories (**Fig. 3b**). We observed an unbinding event in one of the R50S+R59N trajectories (**Fig. 3c**); in other replicas, we also noticed deviations in the packing geometries of the trastuzumab VH and VL domains (**Fig. 3d-e**). Together with previous reports on the relationship between VH-VL packing and paratope shapes^17,54,55^, R50S+R59N could perturb the trastuzumab paratope and compromise its interaction with HER2.

**Figure 3.**
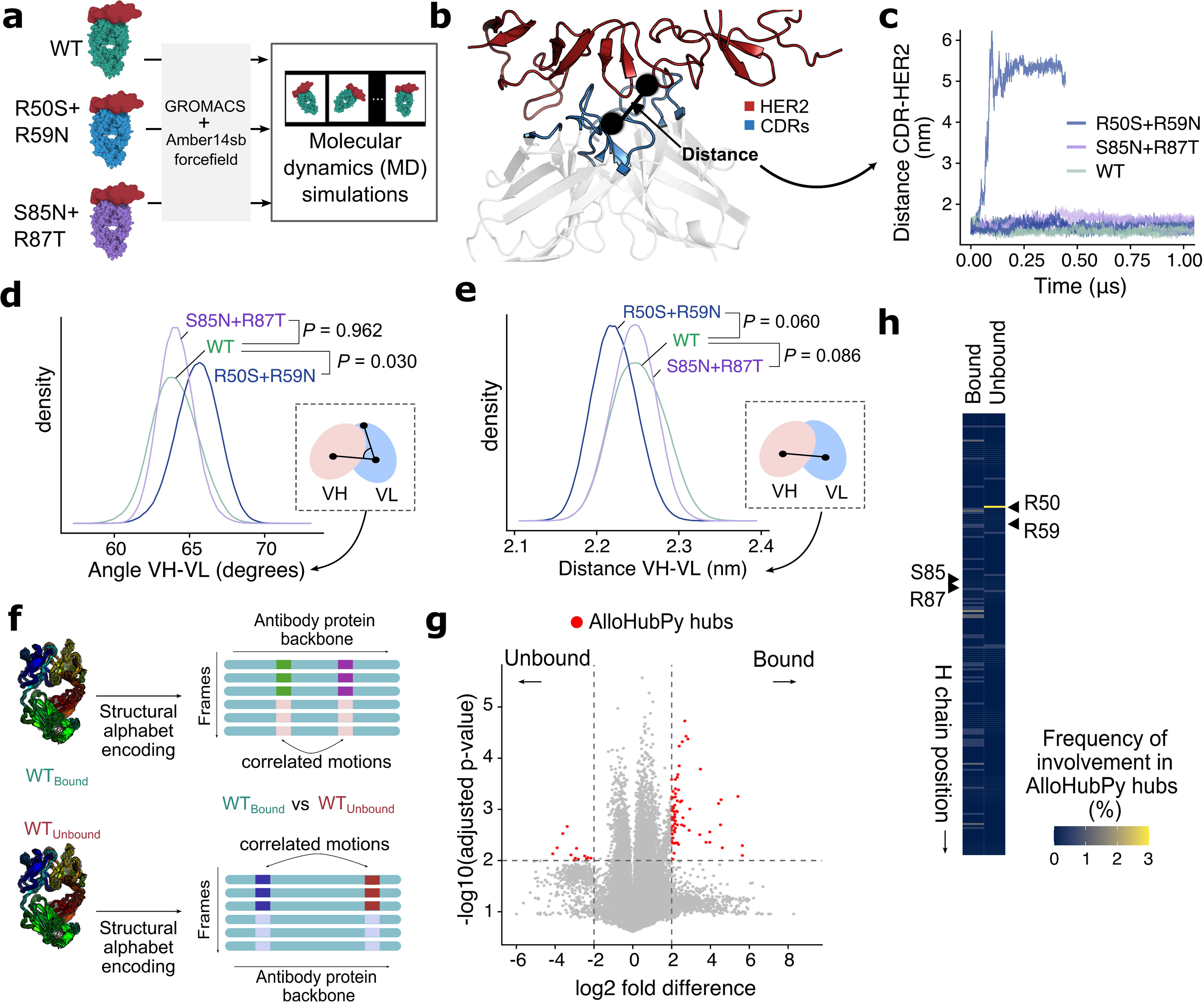
Investigating stabilising trastuzumab VH mutants using molecular dynamics (MD) simulations. (a) Schematic to illustrate the generation of MD simulation trajectories for WT, R50S+R59N and S85N+R87T constructs. (b) Illustration of the calculation of CDR-HER2 distance from MD simulation frames. Black circles depict the centres of mass (COM) of the HER2 juxtamembrane region and the trastuzumab CDR region. (c) Time evolution of CDR-HER2 distance in the MD trajectories. (d-e) Distributions of (d) VH-VL angle (planar angle defined using COMs of the VH, VL and light chain CDR) and (e) VH-VL distance (straight-line distance between COMs of VH and VL domains) over MD simulation frames for the antibody constructs. The inset schematic illustrated the definitions of these measurements. The replica with the HER2 unbinding event (see (b)) for R50S+R59N was excluded from this analysis. P-values were obtained by comparing mutants against WT using linear-mixed effect models. (f) Illustration of analysis of antibody backbone correlated motion using AlloHubPy [ref. 57]. Each MD simulation frame was encoded using the structural alphabet M32K25 [ref. 84], to detect correlated conformational changes between pairwise position (“fragments”) along the antibody structure. This procedure was carried out separately for the WT antibody bound to HER2 (*WT_Bound_*) and that simulated in the absence of HER2 (*WT_Unbound_*) to identify backbone motions which characterised either state. (g) Volcano plot of AlloHubPy fragment pairs whose correlated conformation is found either in the bound (positive log2 fold difference) or the unbound (negative) states. “Hub” positions (absolute log2 fold difference > 2, FDR-adjusted p-values < 0.01) were highlighted. (h) Frequency of AlloHubPy-detected hubs within the trastuzumab heavy chain. Regions corresponding to the sequence position of R50, R59, S85 and R87 were highlighted.

Given the importance of conformational flexibility communicated upon antigen binding from the CDR to more distal sites in the antibody structure^35,36,56^, we asked whether these trastuzumab mutations were associated with local backbone conformational changes important to antigen binding. We utilised our approach (AlloHubPy^57^) to compare MD trajectories of WT trastuzumab in the presence and absence of HER2, to identify positions which displayed significant correlated motions which were associated with HER2 binding (**Fig. 3f-g**, also see Methods). We observed that R50 was located within an AlloHubPy “hub” which displayed significant backbone motions in the unbound state (**Fig. 3h**). This further supported the experimental results that the R50S+R59N mutation could compromise antigen binding.

All together, we have established a pipeline to computationally design novel FW mutations, validate their effects in enhancing antibody stability and function, and integrate MD to interpret mutational impact. This approach identified trastuzumab VH S85N+R87T as a stabilising variant which preserved the capability to bind HER2 and trigger downstream functional effect upon antigen binding.

### Molecular dynamics analysis predicted VL Q89 as a mutable site to tune HER2 binding

We next asked whether we can invert our pipeline, to first use MD to design new mutations guided by conformational dynamics and then validate these designs experimentally. The AlloHubPy analysis of the VL domain revealed one communication “hub” which was present only in the antigen-bound state (**Fig. 4a**). This fragment, Q^89^QHT^92^, extends into the CDRL3 loop. Inspection of trastuzumab structures suggested that the first two glutamine (Q) residues were not exposed at the HER2-CDR interface (**Fig. 4b**). Analysing the preservation of native contacts in the MD simulations of the WT setup, we observed that Q89, but not Q90, retained the majority of its contacts in both the bound and unbound conformational ensembles (Supplementary Figure S7). We therefore explored mutations at the Q89 site to investigate whether this position, despite its lack of direct involvement in HER2 binding, could be tuned to modulate trastuzumab stability and function. We observed in static structures that Q89 formed interactions with surrounding residues in the light chain, including F98 (thus forming the basis of the CDRL3 loop) and Y36 buried in the immunoglobulin fold (**Fig. 4b**). We therefore designed two mutants, Q89H (where the histidine should maintain the local contact network and thus be stabilising) and Q89A (which would introduce flexibility and destabilise local contacts). We generated additional MD trajectories of VL Q89H and Q89A (Supplementary Table S2), and found that Q89H gained stable π–π interaction with F98 (**Fig. 4c-d**). Q89A, on the other hand, lost the majority of local side-chain interactions (**Fig. 4c**), including the contact with F98 (**Fig. 4e**). We expressed Q89A and Q89H *in vitro*, and observed that Q89H maintained a similar thermostability as WT, but Q89A conferred stabilising effects (**Fig. 4f)**. We characterised these antibodies in terms of their capability to bind HER2 (**Fig. 4g**), inhibit HER2+ cancer cell proliferation (**Fig. 4h**) and induce ADCC (**Fig. 4i**). We found that Q89H maintained a similar functionality as WT. Interestingly, despite maintaining a similar capability to bind HER2, Q89A had compromised inhibitory effects to both cell proliferation and ADCC induction (**Fig. 4g-i**). In other words, Q89A was a mutation which decoupled antigen binding from downstream effector function of trastuzumab. We asked whether the MD trajectories provide further molecular details to explain the effects of Q89A, by comparing the contact network between the WT and the Q89 mutants. In contrast with Q89H, Q89A displayed wider perturbation of positions in this network, extending from the CDR region to the VH-VL interface, and propagating to the trastuzumab constant domains (**Fig. 4j**). These analyses highlighted the distal effects of a single mutant and potentially explained the loss of functionality in Q89A that we observed experimentally.

**Figure 4.**
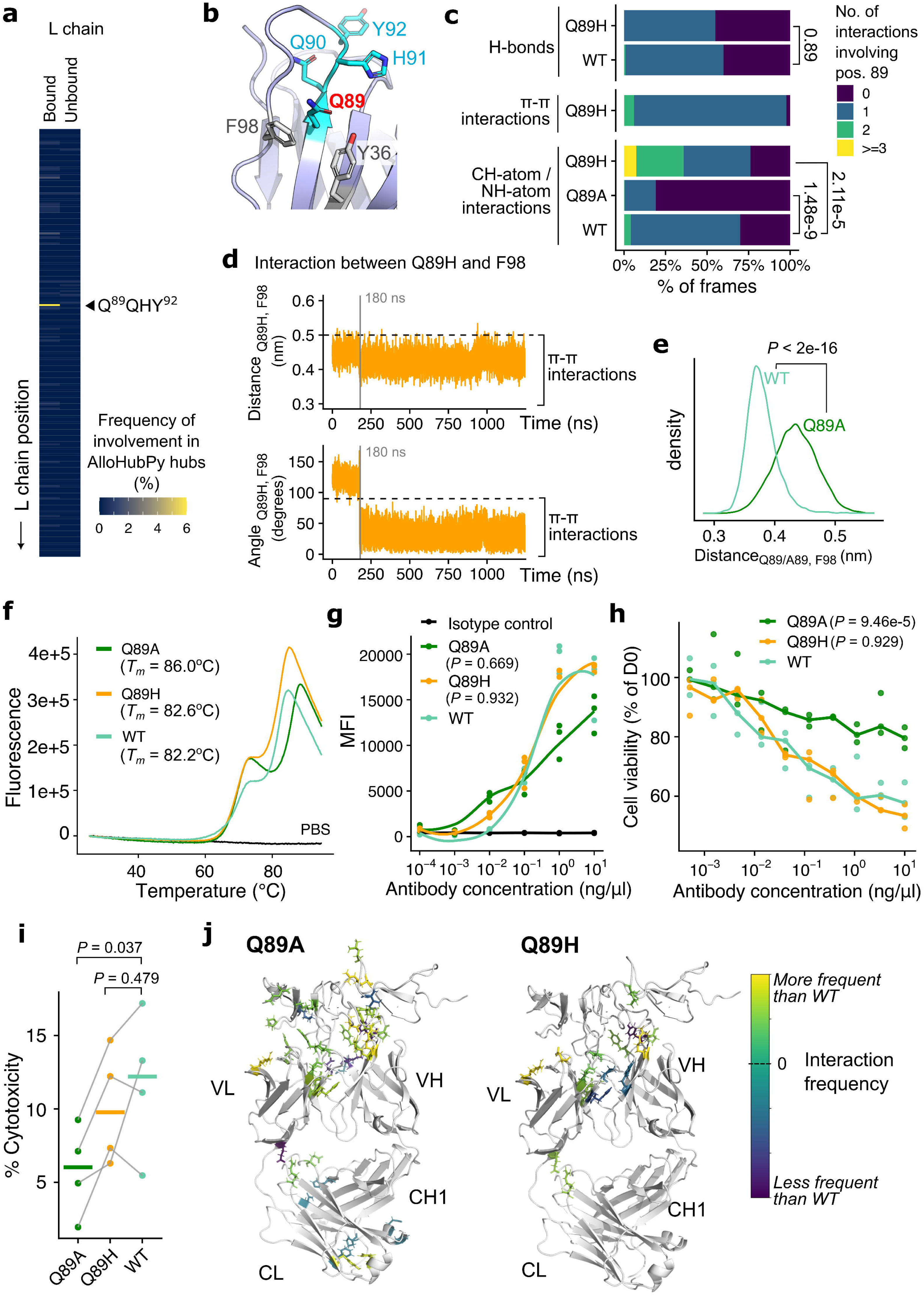
Local backbone dynamics guided the design of trastuzumab VL Q89A to decouple antigen-binding from antibody effector functions. (a) Frequency of AlloHubPy-detected hubs within the trastuzumab light chain. The fragment Q^89^QHY^92^ was frequently observed and highlighted on the plot. (b) Mapping of the Q^89^QHY^92^ fragment on the trastuzumab VL structure. Neighbouring amino acid contacts (F98, Y36) were highlighted. (c) Number of side-chain interactions mediated by WT (Q89), Q89A and Q89H mutants in the MD simulations. Interactions were quantified separately for hydrogen bonds (H-bonds), π – π and CH-atom/NH-atom interactions (see Materials and Methods). P-values were obtained by comparing the distributions of the mutants against the WT using linear mixed-effects models. (d) Time evolution of the (*top*) distance and (*bottom*) angle between Q89H and F98 in the MD simulation trajectory. Dotted lines indicate reference values, below which a π – π interaction occurs. (e) Distributions of the distance between F98 and Q89/A89 in the WT and the Q89A MD simulations. P-value was obtained using a Wilcoxon rank-sum test. (f) Melting curves of WT, Q89A and Q89H determined using differential scanning fluorimetry. Melting temperatures (*T_m_*) of these constructs were noted. (g) Binding of antibody constructs against HER-2 expressing SKBR3 cell lines quantified using FACS. Statistical significance was evaluated using ANOVA followed by post-hoc Dunnett’s tests comparing the mutants against WT. (h) Effect of antibody constructs on SKBR3 cell viability *in vitro*. Statistical significance was assessed with ANOVA followed by post-hoc Dunnett’s tests comparing the mutants against WT. (i) ADCC effects (measured using LDH release as a proxy) on SKBR3 cells co-cultured with NK cells from *n* = 4 donors. P-values were quantified using paired t-tests and displayed after false discovery rate correction. (j) Frequency of side-chain interactions (highlighted with stick representation) observed in the Q89A and Q89H MD simulation frames, compared against the WT trajectories.

## Discussion

Here we established an integrative pipeline combining computational analyses of antibody sequences, static antibody structures, MD simulations and *in vitro* physicochemical and functional assays to design novel FW mutations which modulate antibody stability and function. Antibody FW does not directly engage the antigen, lacks conformational flexibility, and has low sequence diversity^17^. The role of FW in antibody function has thus been underappreciated and less extensively studied in comparison to the CDR regions. Previously, phage display of exhaustive libraries of mutant antibodies have highlighted FW mutations which improve antigen binding and thermostability properties^39^. Here we showed this *in silico* using a structure-based approach (Rosetta) to predict stabilising amino acid substitutions. Forming the bulk of the antibody variable domain, solvent-exposed positions in the FW are inextricably linked to issues with antibody solubility, aggregation, and immunogenicity^20,21,23,31,58^. Given the attention in the field of antibody discovery towards these developability issues, FW mutagenesis is a promising avenue towards optimising antibody designs to become safe and efficacious therapeutics that can be readily manufactured at a large scale. Our work demonstrates that FW mutagenesis can subtly fine-tune antibody properties, not limited to its stability, but also including its antigen affinity and Fab-/Fc-mediated functions. This approach can significantly expand the scope and possibilities of antibody design, instead of restricting design efforts to the CDR regions.

We identified VH S85N+R87T as a stabilising mutant which retained the downstream effector function of trastuzumab, and highlighted the maintenance of packing geometries of the antibody upon mutating S85N+R87T. However, we also noted that these substitutions introduced a novel N-linked glycosylation site. Given the importance of glycosylation in modulating antibody function, we could not exclude the possibility that some of the observed effects of this double mutant may be due to ectopic glycans attached to this site. To further improve and incorporate these considerations, recent approaches in *de novo* protein designs which combine signals from protein structures and sequences^59,60^ may hold promise to optimise design of mutations involving these sites in the future.

A major novelty in this work is the design of a VL mutation (Q89A) which decoupled antigen binding from function: trastuzumab VL Q89A challenges the theory where a high antigen-binding affinity is usually considered as a sign of a functional and efficacious antibody^42,61,62^. Here, this mutation retained antigen-binding, but abrogated the antibody’s ability to inhibit HER2+ cancer cell proliferation and induce ADCC. To the best of our knowledge, this is the first example of such mutation which allows us to dissect the contribution of different structural elements of the antibody to its functional attributes. Our MD analysis highlighted a substantial loss of local contacts in Q89A, extending beyond the immediate vicinity of the mutated residue throughout the variable region FW and beyond. Given previous reports on how CDR-distal mutations affect antibody yield and antigen-binding^18,34,40,62^, our mutagenesis work provided further molecular insights into how the conformational changes upon antigen-binding may be propagated across the entire antibody molecule. Elucidating such mechanisms require a holistic, integrative modelling approach to further probe the importance of different antibody fragments in an antibody’s dynamics^33,41^, specifically on the Fc region, the binding of which to its cognate Fc receptors would induce antibody effector functions^27,36,45^.

An interesting finding is the lack of insights provided by antibody language models (AbLMs) in FW mutagenesis. AbLMs have received widespread attention in the field, due to their ability to generate promising antigen binders using only sequence data^5,12,49^. Further improvements of these models were proposed, including the minimisation of bias towards frequently-used germline immunoglobulin gene fragments^49^, especially given that AbLMs are typically trained from antibody repertoire sampled from peripheral B cells, most of which are of the naïve immunophenotype^63^. We showed here that amino acid likelihoods returned by AbLMs highlighted substantial certainty in antibody FWs, with the wild-type amino acids typically returning a likelihood of 1. This was perhaps unsurprising given the lack of sequence diversity in the FW region in antibody repertoires, leaving the models to focus on the diverse CDR loops in order to learn abstractions from sequence data. For our purpose, these AbLMs on their own thus offered little insights on how FW positions could be optimised further using solely AbLMs. Here we instead adopted a structure-based approach for FW mutagenesis. In the future, dedicated training strategies, for example by generating *in vitro* mutational scanning data specifically on FW substitutions for fine-tuning AbLMs, may help to address this deficiency. This holds promise to streamline the design of FW mutations, and shift the paradigm of antibody design to explore different FW positions in optimising various developability attributes.

In summary, we presented a side-by-side computational-experimental approach to explore the stability and function effects of antibody FW mutations. This approach can potentially be applied beyond antibody FW, for instance to engineer antibody Fc region to enhance specific effector functions, especially given the focus on Fc engineering in recent years for the design of novel antibody therapeutics^27^. Moving forward, considerations beyond the antibody variable domains, including the interrogation of interdomain dynamics^41^, as well as distal effects between the CDR and the Fc, will enable investigations through adopting a holistic view of antibody structural dynamics throughout the entire molecule, and exploring how this can be fine-tuned to achieve desired antibody functions.

## Author contributions

J.C.F.N. conceived the project and planned the investigation under the supervision of F.F. and S.N.K. J.C.F.N. coordinated the bioinformatics analyses, contributed by T.C., D.G., A.K. and D.I. M.L.D.S. performed and analysed the molecular dynamics simulations under the supervision of F.F. and J.C.F.N. A.C. contributed and coordinated the *in vitro* analyses, with contributions from M.G., A.C., A.M., J.C., Y.L. and S.P. J.C.F.N. drafted the manuscript, with input from F.F. and S.N.K. All authors read, commented and approved the final manuscript.

## Acknowledgments

We thank members of the Fraternali and Karagiannis laboratories for helpful discussions, especially to Dr. Oriol Gracia Carmona for his input and to Dr. Silvia Crescioli for initial discussions. The authors acknowledge support by the Biotechnology and Biological Sciences Research Council [BB/T002212/1 to F.F. as principal investigator]; a Career Enhancing Grant from the British Society for Immunology to J.C.F.N.; Worldwide Cancer Research (24-0087); Breast Cancer Now (147; KCL-BCN-Q3) and the CRUK City of London Centre Award (C7893/A29290). D.G. was supported by a studentship from the China Scholarship Council (number 202008440414). A.K. was supported by a Summer Studentship award from King’s College London Faculty of Life Sciences and Medicine. This research was supported by the King’s Health Partners Centre for Translational Medicine. The views expressed are those of the authors and not necessarily those of King’s Health Partners.

## Materials and Methods

### IN SILICO ANALYSIS

#### Antibody sequences and structures

##### Structures

Wherever applicable, for trastuzumab Fab structure we used atomic coordinates from a cryo-EM resolved structure of trastuzumab fragment antigen-binding (Fab) bound to HER2^64^ (Protein Data Bank [PDB] 7mn8). We noted that in this structure, the coordinates for the HER2 juxtamembrane domain, the only portion of the antigen which directly interacts with trastuzumab, were incomplete. For this reason, the input coordinates of HER2 juxtamembrane domain were extracted from PDB 8q6j^65^ (residues 507 to 623 according to author-assigned numbering in the structure), where all the residue coordinates were resolved. We noted that these two structures were highly similar (root mean squared deviation [RMSD] < 1.5Å, calculated on the entire trastuzumab-HER2 complex). Additionally, despite the presence of several glycosylation sites within HER2, none of them is within the juxtamembrane domain. For illustration of structures presented in this paper of trastuzumab in the absence of HER2, a better resolution (PDB 4hkz, resolution = 2.08 Å vs 3.45 Å for PDB 7mn8) structure of trastuzumab^66^ was used.

##### Sequences

For data presented in Figure 1, variable domain sequences of trastuzumab, as well as those of other therapeutic antibodies, were obtained from TheraSAbDab^67^ (accessed May 2024). The list of therapeutic antibodies was further filtered for entries in the format ‘Whole mAb’. This list was overlapped with antibodies with available protein structural data curated in the VCAb database^68^, to obtain structure input for Rosetta. A total of *n* = 55 human antibodies remained for the analysis. Class-switched memory B cell BCR repertoire sequences were obtained from Jaffe et al.^52^. BCR repertoire from intratumoural B cells in breast tumour samples were obtained from Harris et al.^2^. Delineation of CDR and FW followed the IMGT numbering^53^, taken directly from the VCAb^68^ web-tool for sequences extracted from antibody structures and using ANARCI^69^ for sequence data.

### Protein language model analysis

We considered the following protein language models: (i) AntiBERTy^5^; (ii) AbLang2^49^; (iii) AntiBERTa2^48^; (iv) the ‘esm2_t30_150M_UR50D’ version of ESM-2^47^. For each protein language model, we input the heavy and light chain variable region sequences, and tasked the model to predict the amino acid identity of one specified position by masking this position whilst keeping the remaining sequence unmasked. We obtained from each model the output logits for each possible amino acid at the masked position, and converted the logits using the softmax function (in the pytorch package^70^) to probability scores that is stored. This process is repeated for every position in the sequence, resulting in an *n* x *L* matrix, where *n* represents the total number of amino acids (i.e. 20), and *L* is the length of the input sequence.

### *In silico* mutational impact prediction

We predicted the stability effects of mutating every VH and VL positions to every other nineteen amino acids using tools implemented in the Rosetta software suite^71^ (source code version 2018.33.60351 bundle). The trastuzumab Fab coordinates from PDB 7mn8^64^ were subject to backbone conformational sampling using the relax protocol in Rosetta, to generate *n* = 200 structures. Each relaxed structure was subject to the point mutant scan (pmut_scan) protocol for predicting every possible amino acid substitution in the VH and VL regions. The pmut_scan application predicted change in free energy (ΔΔ*G*) which was taken as an indication of whether the mutation was predicted to be stabilising (negative ΔΔ*G*) or destabilsiing (positive ΔΔ*G*). Mutational impact prediction was predicted for double mutants also using the pmut_scan application, with the additional option ‘-double_mutant_scan’ applied when invoking pmut_scan. The ref15 scoring function^50^ was used throughout these protocols.

### Molecular dynamics simulation

Molecular Dynamics (MD) analysis of trastuzumab Fab WT and the mutants studied in this work (VH R50S+R59N, VH S85N+R87T, VL Q89A and VL Q89H) were performed using Gromacs^72^ and the Amber14sb forcefield^73^. The input coordinates for trastuzumab Fab-HER2 complex were extracted from PDB 7mn8^64^, and subjected to superposition with PDB 8q6j^65^ to address missing residues in HER2 in PDB 7mn8. Histidine residues protonation state was assigned through PropKa^74^. All the systems were prepared using an identical procedure detailed as follows: the structure was placed at the centre of a dodecahedron box and filled with TIP3P water. System charge was neutralised by adding counter ions to reach a concentration of NaCl equal to 0.15M. Three minimization steps were then performed, gradually decreasing the heavy atoms’ position restraints. Equilibration runs were started from the minimized structure in the NVT ensemble, gradually increasing the temperature from 50K to 300K. This was followed by NPT equilibration (1 ns) using the Berendsen barostat. Following equilibration, production runs were generated, applying the Parrinello-Rahman barostat to restrain the pressure at 1 bar. The temperature was kept constant at 300K using the velocity-rescale algorithm^75^. Equations of motion were integrated using the Leap-Frog integrator and a timestep of 2 fs. Electrostatic interactions were computed using the Particle Mesh Ewald scheme^76^, with a cutoff of 1.2 nm. The same cutoff value was used for van der Waals interactions.

A table detailing the simulation setups, number of replicates and simulation time can be found in Supplementary Table S2. In brief, a total time of 3 μs was simulated for each system with at least 3 replicates. In R50S+R59N an unbinding event was sampled in the first 0.2 μs of one replicate and not recovered. The simulation was therefore interrupted at 0.5 μs. An additional simulation of the same length was performed for this system. Mutants were produced by changing the selected residue(s) in the input structure using PyMOL (Schrödinger, LLC).

Trajectory analyses were performed using Gromacs tools and the MDTraj^77^ python library (version 1.10.0). π − π stacking was defined on the distance between the centre of mass (COM) of the aromatic rings (hereafter COM_ring_) and the angle between their norm^78,79^. XH-ring (with X=C, N) interaction was determined by examining the distance between the atom X and COM_ring_, as wll as the angle X-H-COM_ring_^80–82^. A detailed list of numeric cut-offs for angles and distances used here to identify molecular interactions can be found in Supplementary Table S3. Hydrogen bonds were assigned according to the Baker-Hubbard criterion^83^. Interaction networks were analysed using the Networkx library (version 3.2.1).

### Identification of local backbone correlated motions upon antigen binding

We used the AlloHubPy^57^ package to analyse our MD simulations for estimating correlated backbone motions within trastuzumab Fab which were associated with antigen binding. We considered the MD trajectories of wild-type trastuzumab Fab in the absence or presence of HER2, using the last 0.5 μs of each replica. Briefly, each frame in the trajectories were coarse-grained at the level of consecutive four-residue backbone fragments using the structural alphabet M32K25^84^. AlloHubPy estimated the normalised mutual information (MI) between each pair of backbone fragments while correcting for systematic errors due to finite conformational sampling^85^. Aiming to identify correlated antibody backbone motions upon HER2 binding, a difference MI matrix was built by subtracting the average MI within MD trajectories of the unbound state from the analogous matrix obtained from trajectories of the bound state. We carried out a differential coupling analysis to identify pairs of fragments (“hubs”) with a log2-fold change in coupling strength of ≥ 2 units and an associated p-value < 0.01.

### IN VITRO ANALYSIS

#### Cells

Human leukocyte cones were obtained from the United Kingdom National Health Service Blood and Transplant system using anonymous donor leukocyte cones following provision of written, informed consent. SKBR3 (ATCC HTB-30) cells were cultured in complete DMEM (10% fetal bovine serum (FBS), penicillin (5,000 U/ml), streptomycin (100 μg/ml)).

### Cell culture conditions

The HER2+ SKBR3 breast cancer model (RRID:CVCL_0033) was obtained from King’s College London Breast Cancer Now Unit, and maintained in high-glucose Dulbecco’s modified Eagle’s Medium (DMEM)-GlutaMAX (ThermoFisher Scientific) with 10% heat-inactivated fetal calf serum. Cells were authenticated by short tandem repeat profiling and used after tested negative for mycoplasma, up to 30 passages. The cells were maintained in a 5% CO2 humidified incubator at 37 °C.

### Flow cytometric analyses

To detect HER2 protein expression levels in vitro, cells were detached with trypsin for 3 min and direct immunofluorescence staining was performed for 20 min on ice using the anti-HER2 IgG1 antibody trastuzumab, followed by a goat anti-human IgG FITC secondary antibody (Vector Laboratories). Samples were acquired using the FACSCanto II flow cytometer equipped with BD FACSDiva Software (BD Biosciences) (RRID:SCR_001456) and data were analysed with FlowJo_V10 software (RRID:SCR_008520) to measure MFI.

### *In vitro* cell viability assay

To measure cell viability, 500 SKBR3 cells per well were plated in 384-well-plates and incubated with treatments for 96 hours at 37 °C. Cell viabilities were detected by CellTiter-Glo Assay (Promega) according to the manufacturer’s instructions. Optical absorbance was read on FLUOstar Omega spectrophotometer (BMG Labtech).

### ADCC assays

Natural Killer (NK) cells were isolated using the RosetteSep^TM^ Human NK Cell Enrichment Cocktail (STEMCELL technologies) following the manufacturer’s instructions. After isolation NK cells were maintained overnight in RPMI1640 supplemented with 10% FBS. The next day NK cells were added to SKBR3 cells at an effector to target cell ratio (E:T) of 10:1 in the presence of antibody (0.01 µg/ml). Samples were incubated for 4 hours at 37°C, 5% CO2. A no antibody control served as spontaneous release control. SKBR3 cells only were lysed 45 min prior to end of the incubation and served as maximum release control. After centrifugation, 50 µL of supernatant were transferred to a clear flat-bottom 96-well plate. Quantification of lactase dehydrogenase (LDH) was performed using CyQUANT LDH Cytotoxicity Assay Kit (Invitrogen) following the manufacturer’s instructions. Absorbance at 490 nm and 680 nm was measured using a FLUOstar® Omega Spectrophotometer (BMG Labtech).

### Differential Scanning fluorimetry

To determine the melting temperatures of the mAbs, 5 mg of each respective mAb in PBS were run in triplicate with SYPRO® Orange (final concentration of 5X) on a QuantStudio7TM Flex Real-Time PCR System. Fluorescence was recorded from 25-95 °C with a ramp of 0.05°C/s. Melting temperatures (*T_m_*) were obtained by determining the local minima of the first derivative of the fluorescence curves.

### STATISTICS AND DATA VISUALISATION

Statistical analysis was carried out in the R statistical computing environment (v4.1.0). Data visualisation was generated with the R ggplot2 package (v3.5.1). Structural visualisation was generated with PyMOL (v3.0.0). For dose-dependent measurements, we fitted an analysis of variance (ANOVA) model where the measured values were modelled as a function of dose and the antibody construct. Unless otherwise stated, for other comparisons we used a linear mixed effect model (using the R lme4 package^86^) incorporating the antibody construct as a fixed effect and replicates as a random effect. Statistical significance was examined via post-hoc Dunnett’s test (R multcomp package), comparing coefficients corresponding to a specified mutant and that of the wild-type. For ADCC assays, statistical significance was evaluated using a paired t-test, with data points paired by biological replicates delineated by the donor origin of the NK cells used in the experiment. For the other data presented herein, statistical significance was assessed using a Wilcoxon rank-sum test.

