## Supplementary Materials for "Tuning antibody stability and function by rational designs of framework mutations"

### Supplementary Tables

Supplementary Table S1. Rosetta saturated mutagenesis results for the VH and VL domains. (XLSX)

Supplementary Table S2. Summary of molecular dynamics simulations.

| System | Number of replicas | Simulation time |
|---|---|---|
| WT with HER2 bound | 3 | 1 μs |
| WT without HER2 (i.e. unbound) | 3 | 1 μs |
| VH R50S+R59N with HER2 | 4 | 1 μs (2 replicas)<br>0.5 μs (2 replicas) |
| VH S85N+R87T with HER2 | 3 | 1 μs |
| VL Q89A with HER2 | 3 | 1 μs |
| VL Q89H with HER2 | 3 | 1 μs |

Supplementary Table S3. Criteria used for accessing residue interactions in MD trajectories. COM, centre of mass.

| Interaction Type | Distance cutoff (nm) | Angle cutoff (degrees) | Notes | Reference |
|---|---|---|---|---|
| π–π stacking | ≤ 0.5 (COM–COM) | ≤ 90 (norm-norm) | Norm defined on the aromatic ring. | [1][2] |
| XH-ring (X = C, N) | ≤ 0.4 ($X–COM_{ring}$) | > 45 ($X-H-COM_{ring}$) | | [3][4] |

### Supplementary Figures

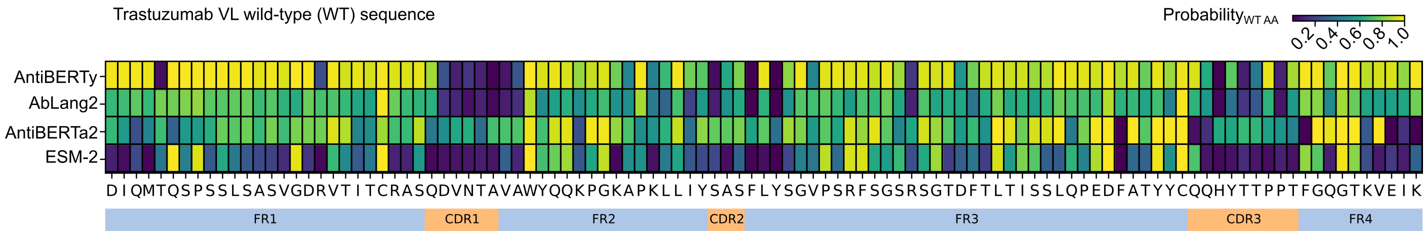


**Supplementary Figure S1**. Heatmap illustrating position-specific probability for the wild-type amino acid (columns) along the trastuzumab VL sequence using different language models (rows). FW and CDR regions are delimited below the heatmap.

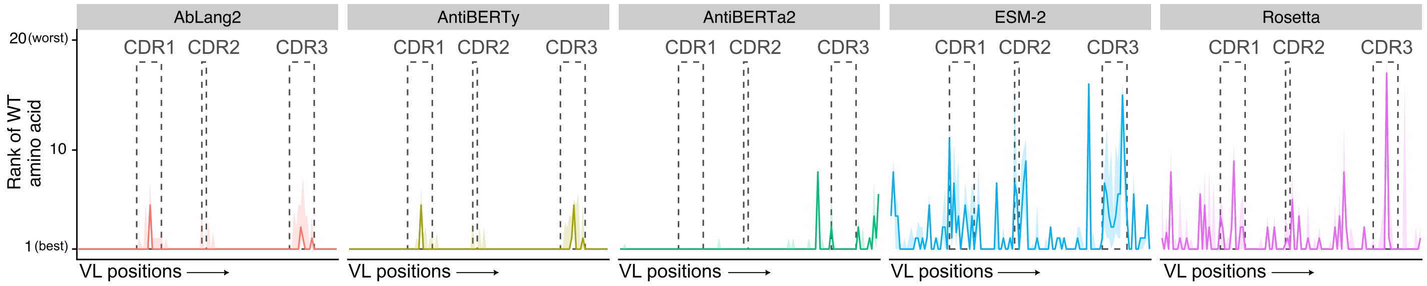


**Supplementary Figure S2**. Rank of wild-type amino acid (vertical axis, 1 = best, 20 = worst) along the VL sequences (horizontal axis) of n = 55 human therapeutic antibodies using the computational approaches evaluated in this work.

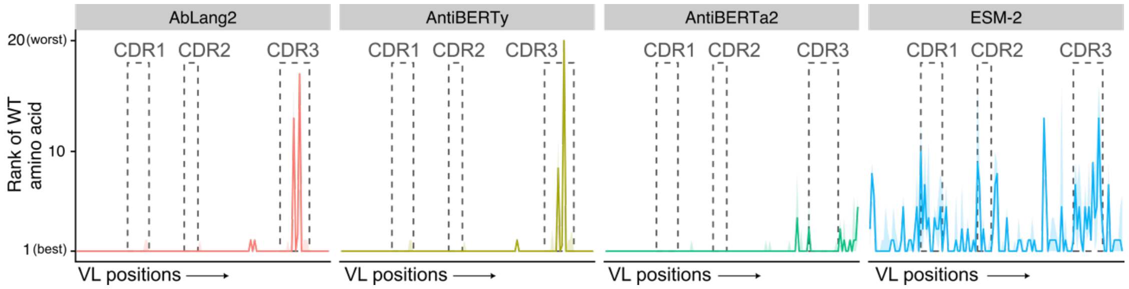


**Supplementary Figure S3**. Rank of wild-type amino acid (vertical axis, 1 = best, 20 = worst) along the VL sequences (horizontal axis) of n = 1,988 paired H-L chains from class-switched memory B cells taken from the Jaffe et al. [5] dataset.

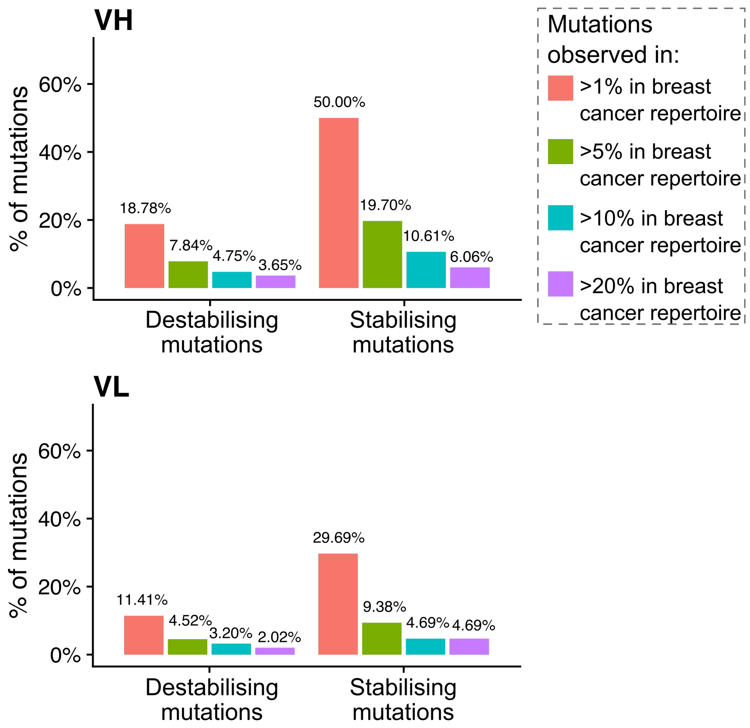


**Supplementary Figure S4**. Comparison of mutation occurrence in a breast cancer antibody repertoire dataset (Harris et al. [6]) against stabilizing mutational effect prediction from Rosetta generated in this study. Mutations were classified into destabilizing (Rosetta predicted $\Delta\Delta G > 0$) and stabilizing ($\Delta\Delta G < 0$), and by their observed frequencies (in terms of percentage of sequences in the repertoire data).

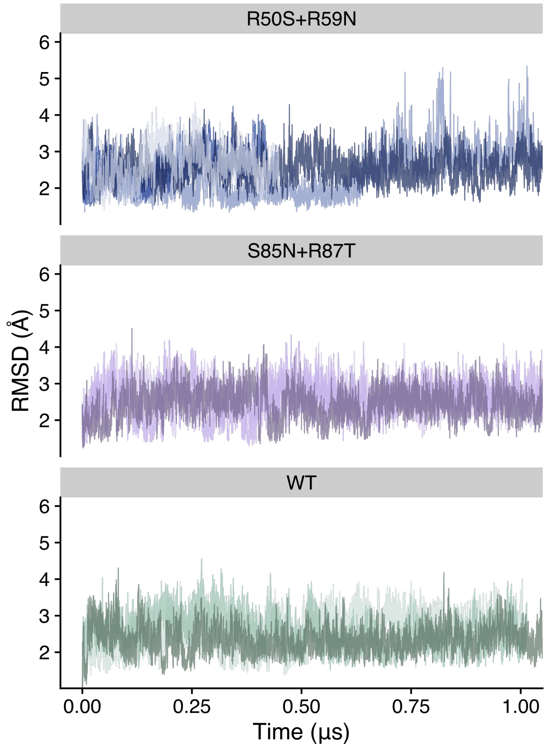


**Supplementary Figure S5**. Root-mean-squared deviation (RMSD) of MD frames across simulation time, for the WT, VH R50S+R59N and VH S85N+R87T systems. Each colour represents a replica.

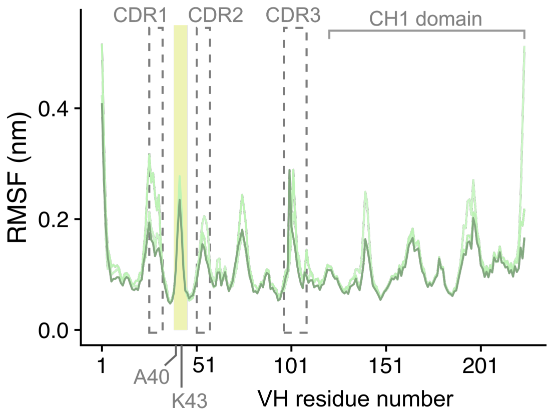


**Supplementary Figure S6**. Root-mean-squared fluctuation (RMSF) of each amino acid Cα in the WT simulations., Each colour represents a replica. The regions corresponding to CDR loops and the CH1 domains are labelled in grey. The loop containing A40 and K43 is highlighted in yellow.

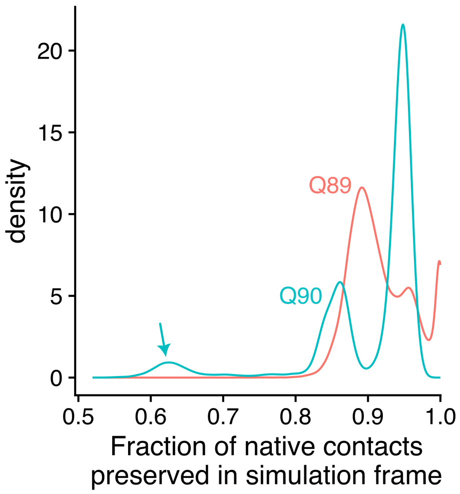


**Supplementary Figure S7**. Fraction of native contacts observed made by Q89 and Q90 in each MD simulation frame in the WT system. 'Native' contacts refer to contacts made by the side-chains of Q89 and Q90, observed in the starting frame and within a shell of 5Å. The subset of frames where Q90 loses native contact was highlighted with an arrow in the plot.
